# Boquila: NGS read simulator to eliminate read nucleotide bias in sequence analysis

**DOI:** 10.1101/2022.03.29.486262

**Authors:** Umit Akkose, Ogun Adebali

## Abstract

Sequence content is heterogeneous throughout genomes. Therefore, Genome-wide NGS reads biased towards specific nucleotide profiles are affected by the genome-wide heterogeneous nucleotide distribution. Boquila generates sequences that mimic the nucleotide profile of true reads, which can be used to correct the nucleotide-based bias of genome-wide distribution of NGS reads. Boquila can be configured to generate reads from only specified regions of the reference genome. It also allows the use of input DNA sequencing to correct the bias due to the copy number variations in the genome. Boquila uses standard file formats for input and output data, and it can be easily integrated into any workflow for high-throughput sequencing applications.

## Introduction

Simulating genomic data for benchmarking bioinformatics programs has become increasingly popular, particularly for read alignment, genome assembly, and variant and RNA-seq analysis [1]. Using such an approach allows for systematic performance assessment even in the absence of gold-standard data. Most currently available simulation tools are heavily geared towards benchmarking; they concentrate on generating reads produced by a specific sequencing experiment by modeling the characteristics of reads produced by sequencing machinery. Consequently, the correction metrics are primarily associated with artificial errors commonly introduced by these specific sequencing protocols.

Although most of the tools use some profile for simulating, these profiles are used for simulating characteristics of sequencing protocols rather than the biological experiments. No simulation tool utilizes nucleotide content profile to our knowledge. SomatoSim [2], VarSim [3], SimuSCoP [4], and many other tools [5–9] were specifically designed to simulate genomic variation. ART [10] and SInC [5] generate profiles based on specific error models and quality score distribution extracted from empirical data. pIRS [11] and Mitty [12] generate quality profiles based on mapped reads and empirical data. NanoSim [13], a nanopore sequence simulator, also uses error profiles and length distributions. Gargammel, ancient DNA sequencing simulator, uses sequencing errors and quality profiles can model base compositions. It can mimic UV damage by adding deamination. However, it is specifically designed for simulation of ancient DNA sequences [14]. BEAR [15], focused on metagenomics, generates error, quality, and abundance profiles. However, the nucleotide content of the reads could be biased for several reasons. First, if sequence library preparation involves immunoprecipitation, antibodies might be biased towards pulling down specific nucleotide profiles. Moreover, ligation efficiencies could be different across varying nucleotides on both 5’ and 3’ ends, which would result in some nucleotide enrichment at the read ends. Furthermore, the nature of sequencing technology might supposedly yield a particular biased nucleotide profile. For instance, sequencing methods yielding the maps of UV damage naturally result in dipyrimidine-enriched reads [16,17]. Additionally, the GC content of the reads might vary depending on the sequencing platform [18]. Finally, the PCR step might introduce another nucleotide bias due to differential efficiencies of universal primers towards specific nucleotides [19]. Considering such factors that could affect the genomic distribution of the reads, there is a clear need for a sequencing read simulation tool that utilizes nucleotide content profile although simulated reads can also be used to eliminate nucleotide content bias in experiments whose results are affected by nucleotide content. Simulation tools mentioned above can account for error and quality profiles of sequencing platforms and GC content biases. However, most of them were designed to simulate reads based on sequencing instruments. There was no other option if aimed to generate a simulated dataset that mimics the nucleotide content of input reads. Here we present boquila, a nucleotide content-based NGS read simulator that can produce simulated reads with a similar nucleotide content profile to input reads. They can be used to normalize the nucleotide content bias in actual reads.

Genomic regions with higher copy numbers have a greater chance of being pulled down during library preparation, whereas those with lower copy numbers are harder to detect. Boquila can also use data from input sequencing as input while generating simulated reads. With this approach, we can also use generated simulated reads to normalize the impacts of copy number variations.

## Features and methods

Boquila was specifically developed to generate synthetic reads with the same nucleotide content as input reads. It operates with FASTA or FASTQ files as an input and generates reads according to the nucleotide content of the input. The number of generated reads and their length distribution will equal the input reads. The nucleotide profile can be calculated based on user-defined kmer length or single nucleotides. Boquila can use the entire genome or pre-defined genomic intervals while randomly selecting reads from the reference genome, thus providing fine-grained control over the regions where simulated reads are generated. Alternative to the reference genome, input sequencing reads can also be used, if a user has raw genome sequence data as a control retrieved from the same experimental setup (cell type, conditions, etc). In this case, reads are randomly generated from input sequencing reads. When generating simulated reads, the nucleotide profile obtained from input reads is adjusted dynamically based on the nucleotide profile obtained so far from simulated reads. In this manner, simulated reads can be further conformed to input reads.

Simulated reads are exported in FASTA or FASTQ format based on the input read format. Simulated reads can also be exported in BED format. Quality scores are copied over from input reads if input reads are present in FASTQ format. FASTA and FASTQ are standard formats for high-throughput sequencing reads, making boquila easier to integrate into existing workflows. Additionally, obtaining output in BED format can help bypass alignment, which is one of the most time- and resource-intensive steps in any NGS workflow.

## Results

We used boquila to generate simulated reads for the data sets of two published studies: XR-seq data from [20] and damage-seq data from [21]. Simulated reads have the same nucleotide content as true reads Fig 1B (XR-seq Hotelling’s T-square: 0.25, p-value: 0.99, damage-seq Hotelling’s T-square: 5.2, p-value: 0.4), but they are randomly generated from the genome. Both XR-seq and damage-seq reads have a certain nucleotide bias due to the UV-induced damage site. The simulated reads generated by boquila can be used to normalize the true repair signal and the true damage signal obtained from XR-seq and damage-seq, respectively Fig 1C. Using this method, we can eliminate the potential bias of the nucleotide content in subsequent analyses.

**Fig 1.**
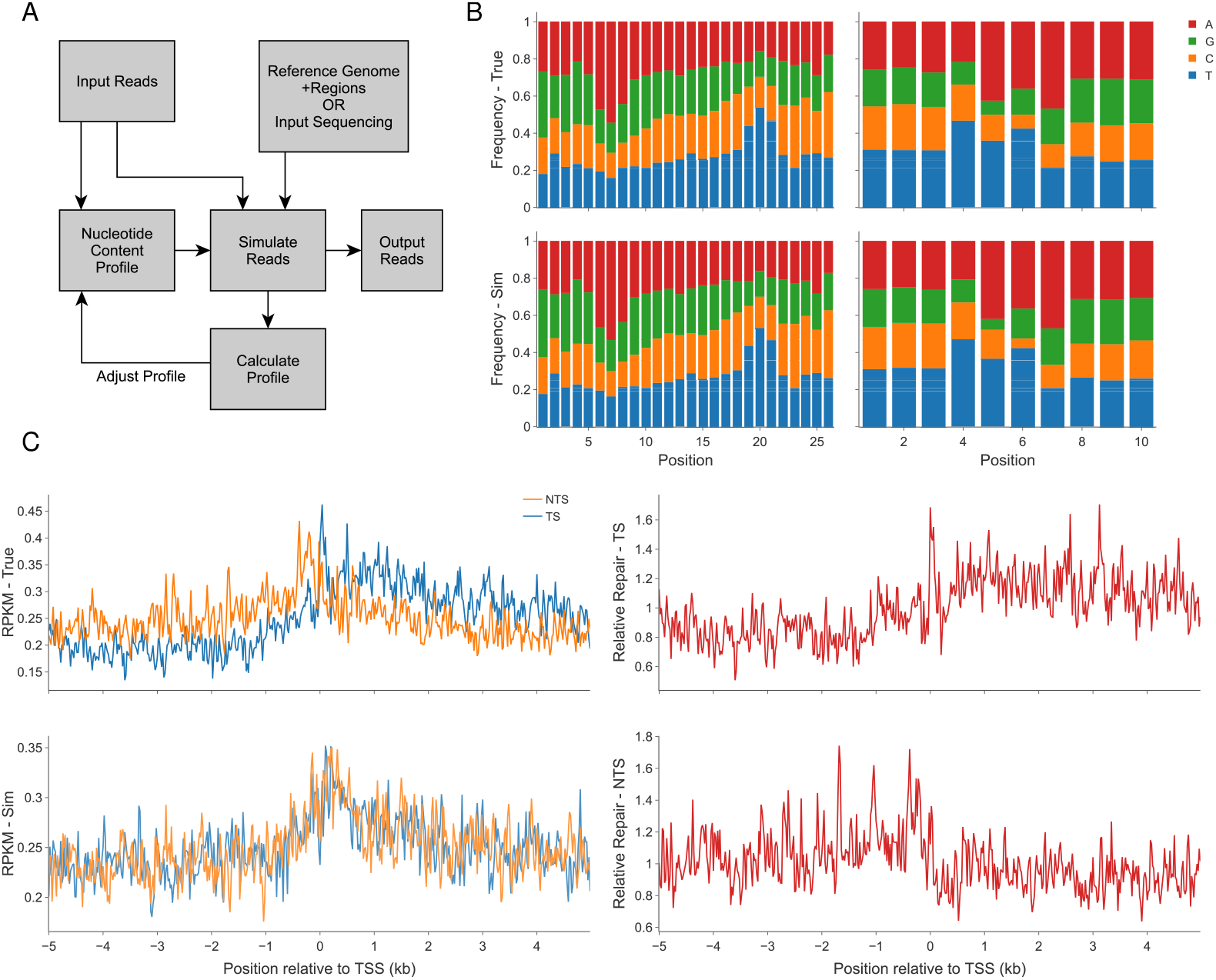
A: Overall workflow of boquila. B: Nucleotide frequency of simulated and true reads for XR-seq (left) and damage-seq (right). XR-seq reads are enriched with thymines at 19-21st positions of the reads, whereas centered damage-seq reads are enriched with pyrimidines at 5-6th positions. C: UV-induced repair (XR-seq) profiles around transcription start sites (TSS). Due to transcription-coupled repair (TCR), transcribed strand (TS) is expected to have higher repair signals relative to the non-transcribed strand (NTS). Whether sequence bias affect the TCR profile around TSS is investigated with the simulated reads (bottom). The profiles showing observed/expected (due to sequence context) ratios (on the right) indicate the TS (top) and NTS (bottom) TCR profiles.

## Discussion

In this study, we propose a novel sequencing simulator utilizing nucleotide content. Some simulators can account for bias produced by sequencing machinery and GC content. Furthermore, some focused on introducing genomic variation into simulated reads, Table 1. However, no simulation tool can yield random reads reflecting the nucleotide content profile of the input reads. Alternative to the reference genome, boquila can also use input sequencing data during simulation to account for CNV bias, which most other simulation tools neglect. With the development of NGS technologies for each specific research question, specialized correction tools must be used to remove technical artifacts of the methodology. In that respect, boquila fills an essential gap in NGS analysis.

**Table 1.** Brief summary of existing simulation tools.

| Simulator | Output Layout | Output Format | GC bias | CNV bias | Nucleotide Content | Genomic Variation | Simulated Profile |
| --- | --- | --- | --- | --- | --- | --- | --- |
| MetaSim [22] | SE | FA |  |  |  |  | empirical error, genome abundance profile |
| Mason [23] | SE,PE | FA,FQ |  |  |  |  | empirical error and quality profiles |
| BEERS [24] | PE | FA |  |  |  |  | intron signal profile, error profile |
| ART [10] | SE,PE | FQ,SAM |  |  |  |  | empirical length, error, quality profiles |
| GemSIM [25] | SE,PE | FQ,SAM | ✓ |  |  | ✓ | empirical quality and error profiles |
| Grinder [26] | SE,PE | FQ,FA |  | ✓ |  |  | position dependent error profile |
| pIRS [11] | PE | FQ | ✓ |  |  | ✓ | empirical base-calling, gc%-depth profiles |
| Wessim [27] | SE,PE | FQ,SAM | ✓ |  |  |  | fragment generation models |
| NeSSM [28] | SE,PE | FQ | ✓ |  |  |  | error and coverage profiles |
| xs [29] | SE,PE | FQ |  |  |  |  | read-length and quality profiles |
| SInC [5] | SE,PE | FQ |  |  |  | ✓ | quality and error profiles |
| CuReSim [30] | SE | FQ | ✓ |  |  | ✓ | error profile |
| FASTQSim [31] | SE | FQ |  |  |  |  | read-length, quality and error profiles |
| BEAR [15] | SE,PE | FQ |  |  |  |  | empirical length, error, quality profiles |
| VarSim [3] |  | FQ |  |  |  | ✓ | mutation profile |
| SCNVSim [6] | SE,PE | FA,VCF |  |  |  | SV,CNV | - |
| IntSim [7] | SE,PE | FQ | ✓ |  |  | ✓ | read-length, quality and error profiles |
| tHapMix [8] |  | BAM |  |  |  | ✓ | error profile |
| gargammel [14] | SE,PE | FQ | ✓ |  |  |  | damage, fragmentation, contamination profiles |
| NEAT [32] | SE,PE | FQ | ✓ |  |  | ✓ | read-length, quality and error profiles |
| NanoSim [13] | SE | FQ |  |  |  |  | error profile, length distribution profile |
| Pysim-sv [9] | SE,PE | FQ | ✓ |  |  | ✓ | - |
| Xome-Blender [33] | PE | BAM |  |  |  | ✓ | - |
| Mitty [12] | SE,PE | FQ, VCF | ✓ |  |  | ✓ | empirical quality and length profiles |
| PaSS [34] | SE,PE | FQ |  |  |  |  | read-length and error profiles |
| SimuSCoP [4] | SE,PE | FQ | ✓ |  |  | ✓ | quality and error profiles |
| PBSIM2 [35] | SE | FQ |  |  |  |  | quality and error profiles |
| SomatoSim [2] | PE | SAM,BAM |  |  |  | SNV | error profile |
| boquila | SE | FQ,FA,BED | ✓ | ✓ | ✓ |  | nucleotide content profile |

SE: single end, PE: paired end, FA: fasta, FQ: fastq, SNV: single nucleotide variation, CNV: copy number variation, SV: structural variation.

## Availability

Boquila is written in Rust and freely available at https://github.com/CompGenomeLab/boquila.

## Funding

This publication has been produced benefitting from the 2232 International Fellowship for Outstanding Researchers Program of TUBITAK (Project No: 118C320 to OA). UA and OA are supported by EMBO Installation Grant (Grant No: 4163 to OA) funded by TUBITAK.

## Notes

### Competing Interest Statement

The authors have declared no competing interest.

